# Post-transcriptional mechanisms distinguish human and chimp forebrain progenitor cells

**DOI:** 10.1101/582197

**Authors:** Daniela A Grassi, Per Ludvik Brattås, Jeovanis G Valdés, Melinda Rezeli, Marie E Jönsson, Sara Nolbrant, Malin Parmar, György Marko-Varga, Johan Jakobsson

## Abstract

The forebrain has expanded in size and complexity during hominoid evolution. The contribution of post-transcriptional control of gene expression to this process is unclear. Using in-depth proteomics in combination with bulk and single-cell RNA sequencing, we analyzed protein and RNA levels of almost 5,000 genes in human and chimpanzee forebrain neural progenitor cells. We found that species differences in protein expression level was often independent of RNA levels, and more frequent than transcriptomic differences. Low-abundant proteins were more likely to show species-specific expression levels, while proteins expressed at a high level appeared to have evolved under stricter constraints. Our study implicates a previously unappreciated broad and important role for post-transcriptional regulatory mechanisms in the evolution of the human forebrain.

## Introduction

Since the evolutionary split between the human and chimpanzee lineages, the human forebrain has evolved to be larger and more complex. These changes, which include both an increased number of cells and differences in cell types and connectivity, are thought to underlie human-specific cognitive functions [1–4]. Despite the importance of these features, there are major knowledge gaps regarding the molecular mechanisms that underlie these evolutionary changes.

Several genetic principles are likely to have contributed to the evolution of the human forebrain. For example, recent studies have provided functional evidence that the emergence of new genes through gene duplication, including SRGAP2, ARHGAP11, NOTCH2NL and TBC1D3, contribute to human specific characteristics of forebrain development [5–9].

In addition, changes at the level of the coding sequence can also influence human evolution [10]. However, the protein-coding regions of the genomes are largely conserved between human and chimp. It has therefore been postulated that divergence in *cis*-acting sequences, which lead to transcriptional changes, are major contributors to the phenotypic differences between the two species [11, 12]. In line with this, epigenomic and transcriptomic analyses have revealed that the human and chimpanzee genomes display widespread differences in enhancer composition as well as transcript abundance in many cell types and tissues, including the brain [13–19] and mechanistic studies of e.g. the FZD8-enhancer have provided functional evidence that differences in the expression level of a gene can contribute to phenotypic differences relevant for human and chimpanzee forebrain development [13].

In addition to *ci*s-acting mechanisms, *trans*-acting effects could also impact on the evolutionary divergence of the human and chimpanzee forebrain [20]. For example, post-transcriptional control by species-specific non-coding RNAs may affect the protein translation from transcripts without affecting transcript levels. The presence of such mechanisms would not emerge from transcript level analysis alone but would require simultaneous protein expression analysis. To date, little is known regarding differences in protein levels when comparing human and chimpanzee forebrain development, and how such differences may relate to altered transcription. Thus, the relative contribution of *cis*- and *trans*-acting effects to human brain evolution remains unclear.

In this study, we explored transcriptional and proteomic differences between chimpanzee and human forebrain neural progenitor cells (fbNPCs). We developed a robust 2D *in-vitro* differentiation protocol allowing for large numbers of fbNPCs to be generated from induced pluripotent stem cells (iPSCs) of human and chimpanzee origin. Using bulk RNA-seq, single-cell RNA-seq, and in-depth proteomics we compared RNA and protein levels for nearly 5,000 genes in fbNPCs. We found that differences in protein expression levels was much more abundant than the differences in RNA levels, and that the majority of changes in protein levels were independent of alterations in RNA levels. These results imply that post-transcriptional mechanisms have had an important and previously unappreciated role in the evolution of the developing human forebrain.

## Results

### Derivation of human and chimpanzee forebrain neural progenitor cells

Comparative transcriptomics and proteomics on chimpanzees and humans have been hampered due the limited availability of material from the developing forebrain of these species, as well as tissue heterogeneity. Recently, iPSCs from chimpanzee and other hominoids have become available [21–23]. These cells are similar to human pluripotent stem cells, and like their human counterparts they can be used to generate many different cell types. This allows for direct comparison between the transcriptome and proteome of chimpanzee and human stem cell derived cultures.

In order to directly compare human and chimpanzee fbNPCs, we optimized a fully defined, feeder-free, 2D differentiation protocol based on dual-SMAD inhibition (Fig 1A). Chimpanzee iPSCs could be maintained *in vitro* under identical conditions to human iPSCs. Upon differentiation of chimpanzee iPSCs, we observed a rapid morphological switch to fbNPC-like morphology, which was morphologically indistinguishable from that of human iPSC-generated fbNPCs (Fig 1B). After two weeks of differentiation, we used immunocytochemistry (ICC) to determine that both human and chimpanzee fbNPCs expressed homogeneous levels of FOXG1 (Fig 1B), a key forebrain marker, while the pluripotency marker NANOG was absent from the cultures (Extended Data Fig. 1A). These results were consistent in two independent human iPSC lines generated by mRNA transfection (RBRC-HPS0328 606A1 and RBRC-HPS0360 648A1, both from RIKEN; from here on referred to as HS1 and HS2, respectively [24]. In addition, two chimpanzee iPSC lines showed similar results: one generated by mRNA transfection (Sandra A, herein referred to as PT1; [23]) and the other with viral vector transduction (PR00818 PTCL-5, herein referred to as PT2; [22]).

**Fig 1.**
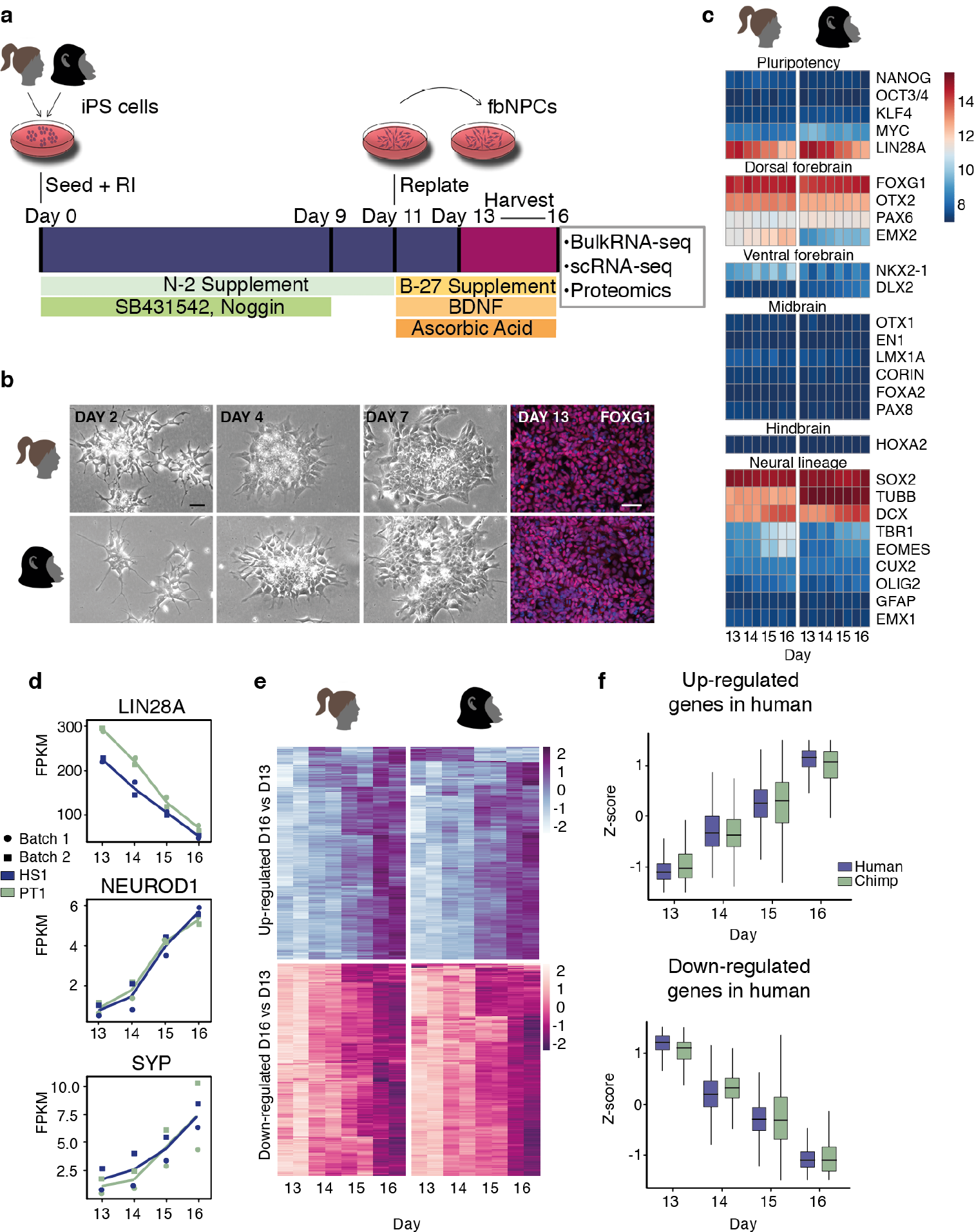
Derivation of human and chimpanzee forebrain neural progenitor cells and temporal progression during forebrain differentiation. (A) Schematics illustrating the differentiation procedure from seeding iPSCs at day 0 to harvesting the forebrain progenitors at days 13-16 of differentiation. (B) Brightfield images of human and chimpanzee cells during the first week of differentiation, and FOXG1 immunocytochemistry at day 13, scale bar represents 50 μm. (C) Heatmap showing marker expression at days 13-16 of differentiation. (D) Dynamic expression of LIN28A, NEUROD1, and SYP. (E) Heatmaps displaying genes that are significantly (p adj. < 0.01) up- and down-regulated over time of differentiation (day 16/day 13) in humans, the same set of genes mapped for both species. (F) Boxplots showing the same set of genes as in E. The lower and upper hinges correspond to the first and third quartiles.

### Human and chimpanzee iPSCs differentiate into forebrain progenitors in a similar temporal progression

To investigate if human and chimpanzee iPSCs would differentiate into fbNPCs with different temporal trajectories, e.g. due to potential differences in cell-cycle progression [23], we performed RNA-seq at 13, 14, 15, and 16 days of differentiation, and analyzed covariance of gene expression between these days and the different species. We mapped the RNA-seq reads from human and chimpanzee samples to the human (GRCh38) and chimpanzee (PanTro6) reference genomes, respectively. To quantify gene expression, we used GENCODE (v27) annotation, which was lifted to the PanTro6 genome to quantify chimpanzee samples.

Transcriptome analysis confirmed that both human and chimpanzee fbNPCs express appropriate neuronal and forebrain markers, while genes related to other brain regions or other tissues were not detected (Fig 1C). At the selected time-points, the fbNPCs corresponded to a differentiation stage just prior to neuronal commitment, demonstrated by the gradual increase in neuronal markers such as *NEUROD1* and *SYP*, and the gradual loss of stem-cell markers such as *LIN28A* from day 13 to day 16 (Fig 1D).

Globally, a similar set of genes were up- or down-regulated between day 13 and day 16 in human and chimpanzee fbNPCs, indicating that the temporal dynamics of the protocol was identical between the two species (Fig 1E&F). Biological replicates confirmed a very limited batch-to-batch variation in the differentiation protocol (Fig 1D, Extended Data Fig. 1B) and the results were consistent in the cell lines from different individuals (Extended Data Fig. 1C).

### Single-cell RNA-seq analysis of iPSC-derived forebrain progenitors

In order to comprehensively compare the cells generated *in vitro* from these two species, we performed single-cell RNA-seq analysis to thereby investigate the heterogeneity of human and chimpanzee fbNPC cultures. At day 14 of differentiation we analyzed transcriptomes of 4,355 human and 3,620 chimpanzee fbNPCs. Using PCA analysis we found that some 95% of the cells clustered into one major population for both species. The transcriptional variation within this main population was mainly explained by differences in cell-cycle state rather than cell identity (Fig 2A); after regressing out cell-cycle effects these cells clustered into a dense population inseparable on PC1 and PC2 (Fig 2B).

**Fig 2.**
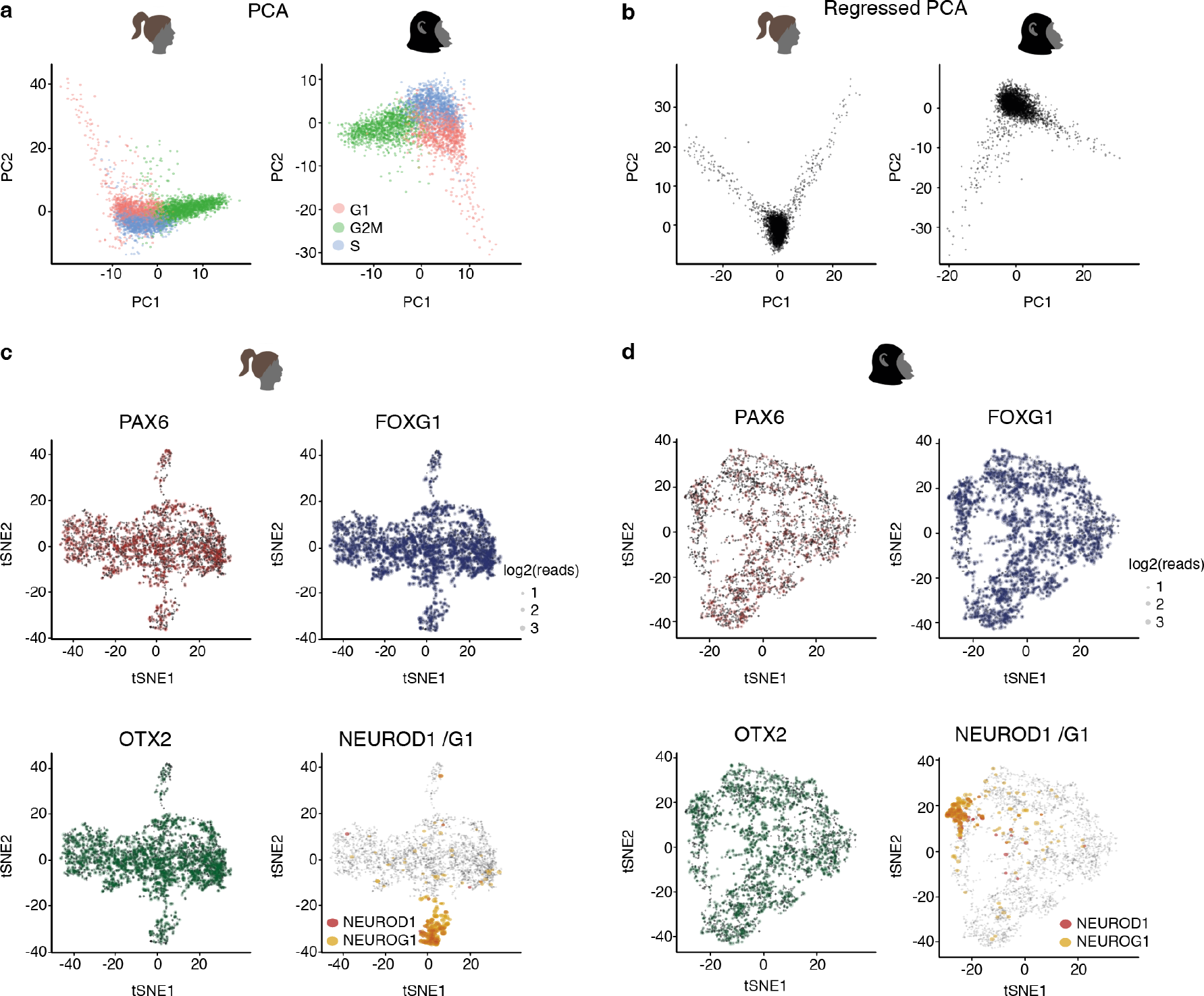
Single-cell RNA-seq analysis of iPSC-derived forebrain progenitors. (A) Principal component analysis of single-cell RNA-seq divided into groups based on cell-cycle stages G1 (red), G2M (green), and S (blue). (B) Principal component analysis of single-cell RNA-seq, where cell-cycle effects have been regressed out. (C-D) Expression of forebrain markers PAX6, FOXG1, and OTX2, as well as neuronal markers NEUROD1/NEUROG1, in human (C) and chimpanzee fbNPCs (D).

tSNE analysis confirmed the presence of a large major population of cells that were homogeneously expressing the forebrain progenitor markers FOXG1, OTX2, and PAX6 (Fig 2C–D). The tSNE analysis also revealed two minor populations, one of which expressed markers associated with early-committed neurons such as NeuroG1 and NeuroD1 (Fig 2C–D, Extended Data Fig. 1D), while the second subpopulation expressed genes related to the endothelial lineage (e.g. ANKRD1 and CTGF; Extended Data Fig. 1D&E). These small subpopulations together made up less than 5% of the cells in both human and chimpanzee fbNPC cultures (Extended Data Fig. 1E).

Taken together, the bulk- and single-cell RNA-seq analysis demonstrate that this 2D differentiation protocol reproducibly gave rise to temporally and phenotypically matched homogeneous cultures of human and chimpanzee fbNPCs, making it a suitable model system for direct comparative analysis.

### Proteomics analysis of human and chimpanzee fbNPCs

To investigate differences between human and chimpanzee fbNPCs at the protein level, we used mass spectrometry (MS) to perform in-depth proteomic analysis on fbNPCs derived from the two human and two chimpanzee iPSC lines (Extended Data Fig. 1F). We differentiated the cell lines in three biological replicates for 14 days. Each biological replicate was run in two MS replicates, which were merged for analysis (Extended Data Fig. 1F). Human and chimpanzee MS output was compared to the human and chimpanzee UniProt reference proteomes, and human and chimpanzee proteins were matched on gene name IDs.

This proteomic analysis resulted in 4,956 proteins being quantified and identified at high confidence (FDR<0.01) in at least two replicates in one cell line, and at least two differentiation replicates in both species. The vast majority of detected proteins represented highly abundant transcripts (Fig 3A). Analysis of selected marker genes revealed that several genes associated with fbNPCs were detected in the proteomic analysis, and at similar levels in the different cell lines from both species (Extended Data Fig. 2A). We also verified the absence of proteins known to be expressed in other brain regions or different tissues (Extended Data Fig. 2A). Comparing protein expression levels of the 4,956 genes between the two individuals of the same species revealed a very high degree of correlation, similar to the correlation found when comparing bulk RNA levels (Pearson correlation of 0.984 and 0.98 for human and chimpanzee RNA, and 0.979 and 0.971 for human and chimpanzee protein). These results demonstrate the technical robustness and reproducibility of this approach (Extended Data Fig. 2 B&C).

**Fig 3.**
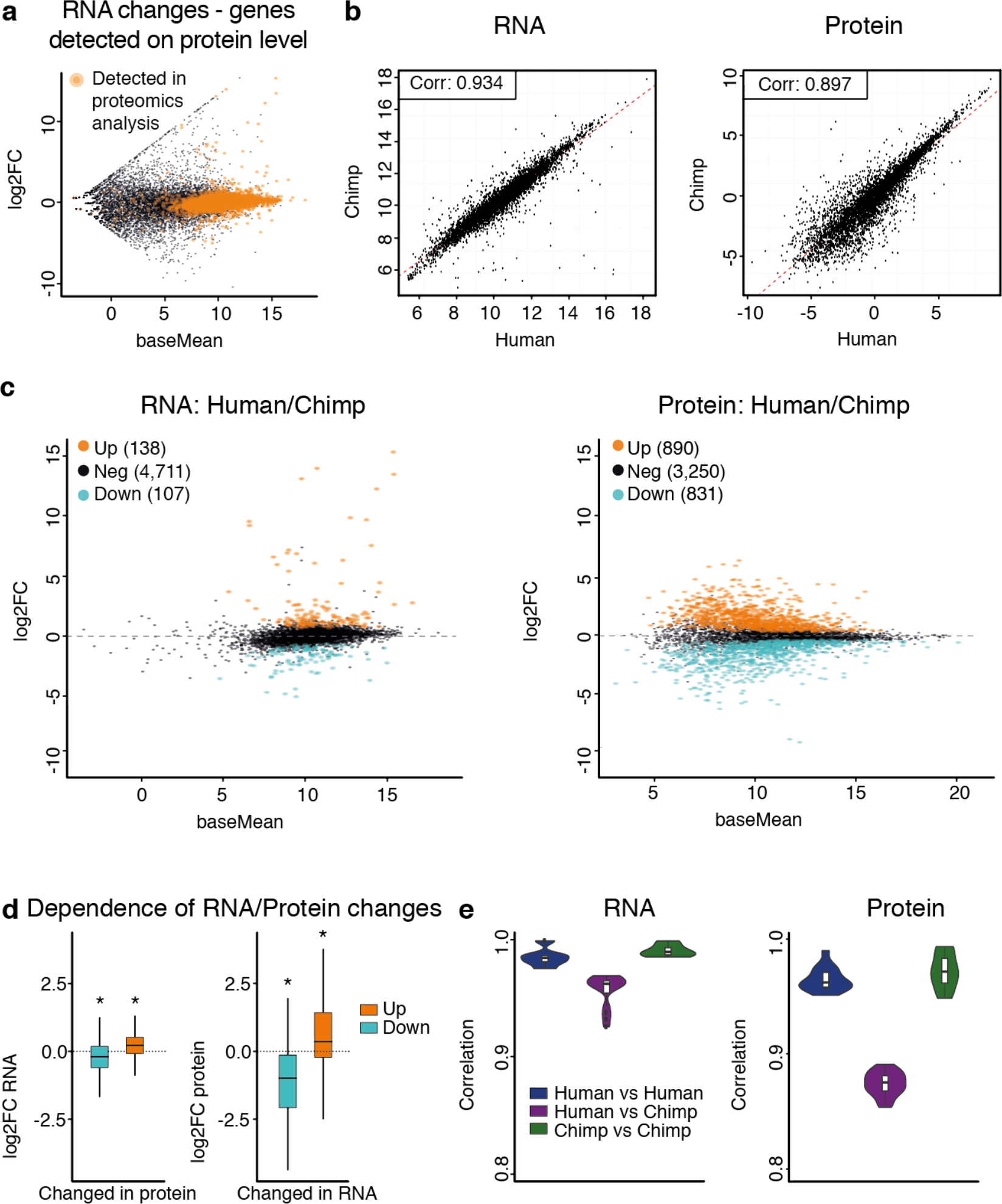
Proteomics analysis of human and chimpanzee fbNPCs. (A) MA plot displaying genes that are detected by proteomics (orange) and genes only detected at the RNA level (black). (B) Human-chimpanzee correlation of RNA abundance, Pearson corr. 0.934, and protein abundance, Pearson corr. 0.897. (C) MA plot of differentially-expressed genes, with 107 down-regulated genes in human compared to chimp, and 138 genes up-regulated at the RNA level, and 831 down-regulated and 890 up-regulated at the protein level. (D) Boxplots showing dependence of RNA and protein changes on differentially-expressed proteins and RNAs. The lower and upper hinges correspond to the first and third quartiles * = p-value = 0.0001, permutation test. (E) Violin plots showing the abundance correlation of RNA and protein between HS1/HS2 (human vs human), human vs chimp, and PT1/PT2 (chimpanzee vs chimp).

### Extensive protein expression differences between human and chimpanzee forebrain neural progenitor cells

In order to investigate how transcriptional differences between human and chimpanzee fbNPCs related to differences in protein levels, we compared differences in RNA and protein levels of the 4,956 genes detected in both RNA-seq and proteomics between human and chimpanzee fbNPCs. We found a strong correlation, at the RNA level, between species regarding expression of these genes (Fig 3B, Pearson correlation: 0.934), and only 245 genes were significantly differentially expressed between human and chimpanzee (Fig 3C, BH-corrected p-value<0.001). Strikingly, there was much less correlation at the protein level (Fig 3B, Pearson correlation: 0.897) where 1,721 genes were differentially expressed between human and chimpanzee fbNPCs (Fig 3C, moderated p-value<0.001). Together, our data suggest that changes at the post-transcriptional level is the major contributor to differences in protein level expression when comparing developing human and chimpanzee forebrain cells.

In order to compare how RNA and protein abundances relate to each other at the single-gene level we first analyzed the RNA levels of genes where the protein level significantly differed between the species. We found that alterations in protein levels relate to RNA levels in a significant but minor way (log2 mean fold change of −0.24 for downregulated and 0.27 for upregulated proteins, Fig 3D). In contrast, when analyzing the protein levels of genes that were differently expressed at the RNA level, we found a much stronger relation between RNA and protein (log2 mean fold change of −1.21 for down- and 0.72 for up-regulated RNAs, Fig 3D), suggesting that most alterations at the RNA level resulted in corresponding changes at the protein level. Taken together, this analysis indicates that there are two principles driving protein expression level differences, when comparing between human and chimpanzee fbNPCs, characterized by the dependence and independence of changes in RNA-level, where the latter contributes to the majority of proteomic differences.

These findings were surprising, since a previous study performed in lymphoblastoid cell lines suggested that post-translational buffering led to the opposite result: higher differences in RNA levels when compared to protein levels [25]. We therefore performed a series of control analyses to corroborate our findings. First, we confirmed that technical variation was not responsible for the larger differences in the proteomics analysis (Extended Data Fig. 2D). We also confirmed that intra-species variation was similar and minimal when investigating either RNA or proteins (Fig 3E). In addition, we also found that the correlation of RNA and protein expression levels in chimpanzee and human fbNPCs were very similar (Extended Data Fig. 2E, Pearson’s correlation: 0.427 in human and 0.447 in chimpanzee) and that this rate of RNA/protein correlation was in line with previous studies (see e.g. [25]). Thus, we concluded that differences in protein levels that is independent of changes in RNA levels, accounted for most of the variation in protein content observed between human and chimpanzee fbNPCs.

### Protein expression differences between human and chimpanzee fbNPCs in low-abundant proteins

We noted that the majority of proteins displaying differential expression between human and chimpanzee appeared to be expressed at low levels (Fig 3D&E). To analyze this further, we divided the 4,956 genes into four subgroups (n = 1239 for each bin) based on their absolute protein expression levels (Fig 4A). We found that most genes that displayed changes at the protein level were found in the *low* or *mid-low* bin (Fig 4A–C), corresponding to proteins expressed at low or intermediate levels, and we observed modest changes for proteins expressed at high levels. In contrast, genes that displayed differences at the RNA level were enriched for proteins that were expressed at high levels (Fig 4B&C). It is worth noting that highly expressed proteins that displayed alterations at the RNA level displayed very minor differences at the protein level, indicative of post-transcriptional buffering to erase species-specific differences in highly abundant proteins, in line with previous observations [25].

**Fig 4.**
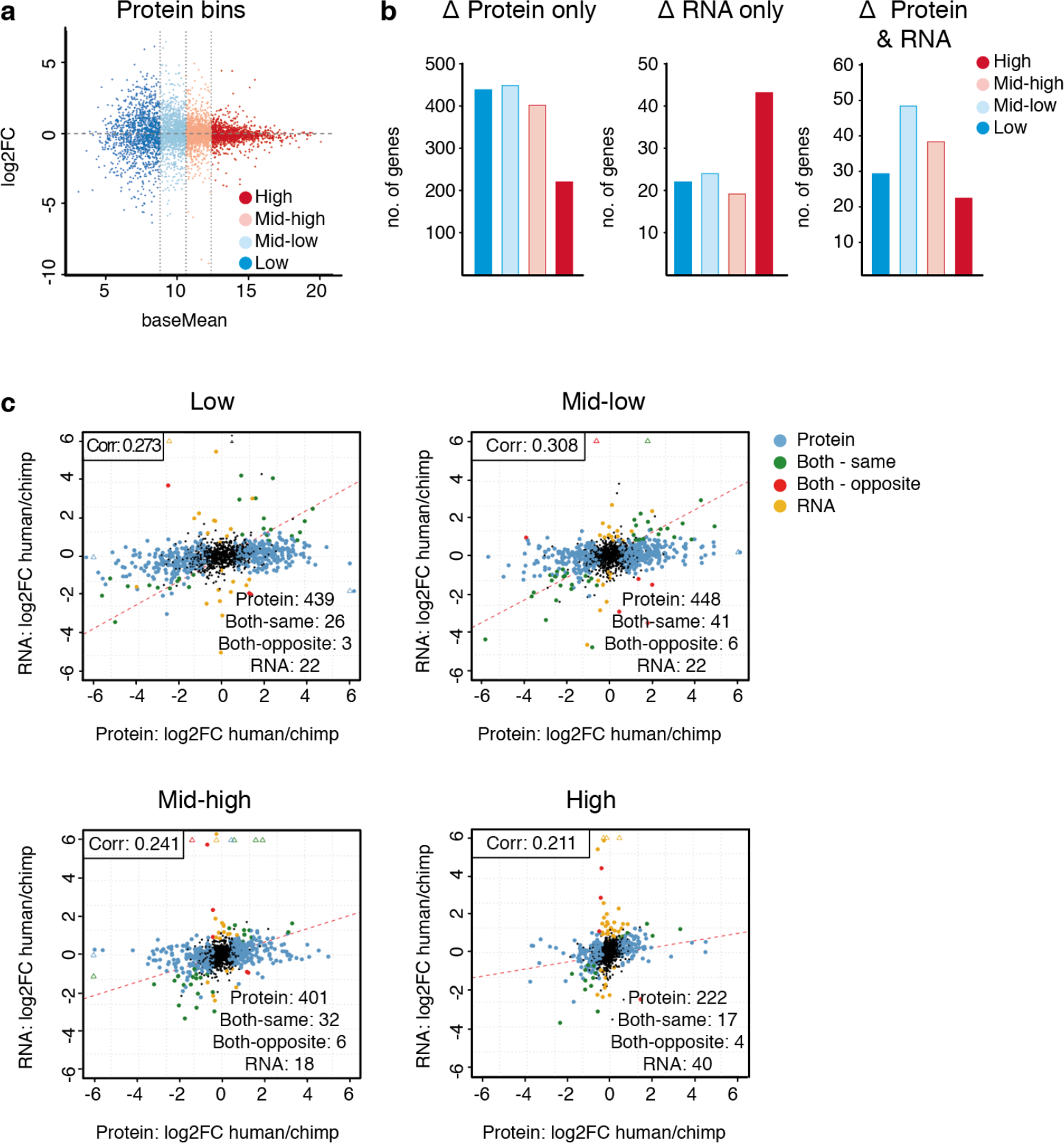
Protein divergence is enriched in proteins expressed at low levels. (A) MA plot showing the log2FC of protein between the two species as a function of protein expression, and the division into 4 subgroups (n = 1239 in each group) based on the level of expression. (B) Bar plots showing the number of proteins and RNAs with a differential expression between humans and chimpanzees at the protein level only, RNA only, and changed in both RNA and protein. (C) Scatter plots showing the RNA and protein fold changes in each of the four subgroups, high, mid-high, mid-low, and low. Genes that are changed at the RNA level only are shown in yellow, protein only in blue, changed in the same direction at both levels in green, and opposite directions in red.

We performed a gene ontology (GO) analysis to investigate if genes displaying alterations at the protein level were enriched for certain biological processes (Extended Data Fig. 3A). We found no positive enrichment for genes that are differentially expressed at the protein level only, while terms involved in RNA processing and metabolic processes were negatively enriched. We also investigated if the number of protein-protein interactions (PPIs) were related to differences in human-chimpanzee protein expression. By integrating StringDB PPI data, we found that proteins with many interactions also displayed a small difference in protein levels between human and chimpanzee fbNPCs, while proteins displaying high differences were enriched for proteins with a low number of PPIs. This was true both for strictly physical binding interactions and for all PPIs (binding, catalysis, reaction, expression, post-translational modification, and inhibition) (Extended Data Fig. 3B). In contrast, the same analysis based on differences in RNA levels showed little difference in the number of PPIs (Extended Data Fig. 3C). Taken together, these data indicated that proteins with different expression between human and chimpanzee fbNPCs are expressed at low levels, not present in large protein complexes, or enriched for particular functional properties.

## Discussion

The study of human and chimpanzee neural tissue has been restricted due to limited availability of material, complex tissue organization, and ethical concerns. Here, we circumvented some of these obstacles by optimizing a 2D culture model of iPSC-derived fbNPCs and generated homogeneous populations of fbNPCs, as evident by the expression of appropriate dorsal forebrain markers. We also found that human and chimpanzee cells differentiated in a remarkably similar temporal fashion and showed limited batch-to-batch variation. Intra-species analysis confirmed that the major human-chimpanzee differences were not due to noise or low sample number. Thus, this culture system offers a robust and feasible model for performing direct comparative analysis between human and chimpanzee forebrain progenitors.

The human forebrain has evolved to be larger and more complex than that of the chimpanzee, even though our genes largely share the same coding sequence [11]. Previous work has pointed to species-specific *cis*-regulatory elements, resulting in diverging transcriptomes, as a contributing factor in brain development [13–19]. However, protein expression levels ultimately determine the phenotype, and the relationship between transcript and protein levels in brain development is not well understood. Here, we performed a direct comparative analysis of protein and RNA levels in human and chimpanzee fbNPCs. Intriguingly, this uncovered higher interspecies expression level differences at the proteomic than the transcriptomic level.

Post-transcriptional control of protein levels can occur in a number of different ways, including mRNA localization, *trans*-acting non-coding RNAs, translational regulation, and selective protein degradation and adds a layer of regulation when transcription is tightly controlled. In fact, post-transcriptional regulation through the binding of RNA-binding proteins has been implicated in several steps of corticogenesis and post-transcriptional regulation of transcription factors is emerging as an important mechanism in controlling neurogenesis [26, 27]. For example, post-transcriptional regulation of Ascl1, a key neurogenic transcription factor, is required for maintenance of adult neural stem cells [28]. It is also worth noting that previous studies on transcriptional differences between several human and chimpanzee tissues have shown lower differential expression in the brain when compared to other organs suggesting that post-transcriptional control may be particular important in the brain. [15, 29]. Our data suggest that the differences found in low abundant proteins contribute to species-specific differences in fbNPCs, and that the regulation of protein expression levels may play a key role in human forebrain development. Our data also support the notion that changes in post-transcriptional and post-translational regulation is a rich source for species-specific divergence in primate forebrain development.

Intriguingly, our results oppose previous data from lymphoblastoid cell lines, where RNA was found to be less tightly regulated, whereas differences in protein expression levels were buffered [25]. Khan et al. [25] reasoned that mRNA expression is under lower evolutionary constraint than protein levels, which could be regulated through post-transcriptional mechanisms. Our data supports the existence of post-transcriptional buffering, but only for RNAs encoding highly-expressed proteins, whereas low abundant proteins were more differentially expressed at the protein level. In the light of this discrepancy between the two studies, it is worth noting that Khan et al. used a different type of quantitative proteomics analysis, with an older instrument that only allowed for the quantification of the 3390 most highly expressed proteins, compared to 4956 in our study. We found that proteins with a conserved expression pattern across the two species were enriched for proteins with a higher number of protein-protein interactions. This indicates, in line with the findings of Khan et al., that the more interactions a protein has, the higher the evolutionary constraint to maintain the same expression level. Additionally, our GO analysis indicates that the expression levels of proteins associated with basic cellular function (e.g. metabolic processes) were more strictly regulated in humans and chimpanzees.

In summary, we have begun to uncover the relationships between transcript and protein expression levels in human and chimpanzee forebrain development. Our results suggest that post-transcriptional and post-translational mechanisms add additional layers of gene regulation, with broad consequences in the developing human brain. Our results provide many new candidate proteins involved in human speciation, and our future research will focus on the function of these genes and the molecular pathways involved in their regulation.

## Materials and Methods

### Experimental design

In this study, we have investigated RNA and protein level differences in human and chimpanzee fbNPCs in order to elucidate the molecular basis for human forebrain development. To this end, iPSCs from two human and two chimpanzee individuals were differentiated into fbNPCs using a 2D culture system with dual SMAD-inhibition. One individual from each species was used to determine potential temporal differences using bulk RNA-seq at day 13, 14, 15, and 16 of differentiation. Day 14 was selected as the time point for further analysis, which included single cell RNA-seq, as well as bulk RNA-seq and in-depth proteomics. Single cell RNA-seq was used to assess the homogeneity of the cultures. Bulk RNA-seq and protein data was compared to determine interspecies expression differences at two levels in forebrain progenitors, as well as determining the relationship between transcript and protein levels for individual genes.

### iPSC culture

iPSCs were maintained on LN521-coated (0.7 μg/cm^2^; Biolamina) Nunc Δ multidishes in iPS brew medium (StemMACS iPS-Brew XF and 0.5% penicillin/streptomycin (Gibco)). Cells were passaged 1:2-1:6 every 2-5 days by being rinsed once with DPBS (Gibco) and dissociated using 0.5 mM EDTA (75 μl/cm^2^; Gibco) at 37°C for 7 minutes. Following incubation, EDTA was carefully aspirated from the well and the cells were washed off from the dish using washing medium (9.5 ml DMEM/F-12 (31330-038; Gibco) and 0.5 ml knockout serum replacement (Gibco)). The cells were then centrifuged at 400 × g for 5 minutes and resuspended in iPS brew medium supplemented with 10 μM Y27632 (Rock inhibitor; Miltenyi) for expansion. The media was changed daily.

### Differentiation into forebrain neural progenitors

iPSCs were grown to a density of approximately 70-90% confluency and were then dissociated as usual for passaging. After centrifugation, the cells were resuspended in N2 medium (1:1 DMEM/F-12 (21331-020; Gibco) and Neurobasal (21103-049; Gibco) supplemented with 1% N2 (Gibco), 2 mM L-glutamine (Gibco), and 0.2% penicillin/streptomycin). The cells were manually counted twice (without the use of trypan blue) and plated at a density of 10000 cells/cm^2^ in 250 μl medium/cm^2^ on LN111 Nunc Δ multidishes (1.14 μg/cm^2^; Biolamina). 10 μM SB431542 (Axon) and 100 ng/ml noggin (Miltenyi) for dual SMAD inhibition, as well as 10 μM Y27632 was added to the medium. The medium was changed every 2-3 days (N2 medium with SB431542 and Noggin) up until day 9 of differentiation, when N2 medium without SMAD inhibitors was used. On day 11, the cells were replated by washing twice with DPBS followed by adding StemPro accutase (75 μl/cm^2^; Gibco) for 10-20 minutes at 37°C. The dissociated cells were washed off with 10 ml wash medium, centrifuged for 5 minutes at 400 × g and resuspended in B27 medium (Neurobasal supplemented with 1% B27 without vitamin A (Gibco), 2 mM L-glutamine and 0.2% penicillin/streptomycin). The cells were counted twice manually and replated at 800,000 cells/cm^2^ on LN111-coated plastic in B27 medium (600 μl medium/cm^2^) supplemented with Y27632 (10 μM), BDNF (20 ng/ml; R&D), and L-ascorbic acid (0.2 mM; Sigma-Aldrich). The cells were kept in the same medium until day 14, after which new B27 medium was added.

### Immunocytochemistry

The cells were washed once with DPBS and fixed for 15 minutes with 4% paraformaldehyde (Merck Millipore), followed by three rinses with DPBS. The fixed cells were then pre-blocked for a minimum of 30 minutes in a blocking solution of KPBS with 0.25% triton-X100 (Fisher Scientific) and 5% donkey serum. The primary antibody (rabbit anti-FOXG1, 1:50 dilution, Abcam, RRID: AB_732415 and anti-NANOG, 1:100 dilution, Abcam, RRID: AB_446437) was added and incubated overnight. On the following day, the cells were washed twice with KPBS and pre-blocked for at least 10 minutes in blocking solution. The secondary antibody (donkey anti-rabbit Cy3; 1:200; Jackson Lab) was added with DAPI (1:1000; Sigma-Aldrich) as a counterstain and incubated at room temperature for one hour, followed by 2-3 rinses with KPBS. The cells were then visualized using a Leica microscope (model DMI6000 B), and images were cropped and adjusted in Adobe Photoshop CC 2015.

### Bulk RNA sequencing

On the day of harvest, the cells were washed once with PBS and lysed with 350 μl RLT buffer with 1% β-mercaptoethanol (Thermo Fisher). The RNA was then extracted using the RNeasy mini kit (Qiagen) according to manufacturer’s protocol. The quality and concentration of the RNA was analyzed using 2100 Bioanalyzer (RNA nano; Agilent) and Qubit (RNA HS assay kit). Libraries for sequencing were prepared using the TruSeq RNA Library Prep kit v2 (Illumina) and again quality-controlled using the Bioanalyzer (high-sensitivity DNA assay) and Qubit (dsDNA HS assay kit). Finally, the libraries were sequenced using an Illumina NextSeq 500, 150× paired-end reads (300 cycles).

Human RNA sequencing samples were mapped to the human reference genome (GRCh38) and chimpanzee samples were mapped to the chimpanzee reference genome (Clint_PTRvs2/PanTro6) using STAR aligner v2.5.0a[30], allowing 0.03 mismatches per base and multimapping at up to 10 loci. Gencode (v27 [31]) gene models were used for splice junction annotation. Gene counts were quantified using the Subread package FeatureCounts [32], counting reads overlapping Gencode (v27) gene annotations. To quantify chimpanzee gene expression, the human Gencode annotation was lifted over to PanTro6 coordinates using the UCSC LiftOver tool. Normalization and differential expression analyses were performed with the R package DESeq2 [33]. All genes annotated as protein-coding in Gencode were used for these calculations. The R package limma was used to correct batch effects [34].

To analyze protein-protein interactions, the protein actions table from StringDB was downloaded ([35], https://stringdb-static.org/download/protein.actions.v10.5.txt.gz, 2018-12-07). Only interactions with score ≥400 were used in the analysis, to exclude low-confidence interactions. To extract only physical binding instances, only interactions with “binding” mode were included from the table. For all interactions, the entire table was included.

Gene ontology enrichment analysis was performed using the Panther Overrepresentation test (Released 20181113) with PANTHER v14.0, released 2018-12-03 [36]. To evaluate significance, all genes detected both in RNA-seq and proteomics were used as the reference background list, to avoid bias from only analyzing a subset of all possible genes. Fisher’s exact test with no correction for multiple testing was used to test for overrepresentation.

### Single-cell RNA sequencing

HS1 and PT1 were differentiated to day 14, washed twice with DPBS, and dissociated with Accutase for approximately 10 minutes, followed by centrifugation at 400 × g for 5 minutes in wash medium. All pipetting was done very gently to avoid cell death. The pellets were resuspended in 1 ml PBS with 0.04% BSA (Sigma) and filtered through 100 μm cell strainers (Falcon) twice. Cells were resuspended in order to yield a concentration of approximately 1000 cells/μl. The single-cell libraries were prepared with Chromium Single Cell A chip kit and Chromium Single Cell 3’ Library & Gel Bead kit v2 (10× Genomics), quality controlled, and quantified using Qubit ds DNA HS and Bioanalyzer High Sensitivity DNA Assay prior to sequencing. The samples were sequenced for 26 cycles on read 1 and 98 cycles on read 2 using the Illumina NextSeq 500.

Raw single-cell RNA-seq data was processed using the Cell Ranger software suite. Raw base call files were converted using cellranger mkfastq before aligning, filtering, barcode count, and UMI counting was performed using cellranger count. Count matrices were further analyzed using the Seurat R package [37]. The data were filtered on number of genes detected in each cell (2000-6000 genes/cell in human and 2000-5000 genes/cell in chimpanzee were kept for further analysis), and only cells with max 0.05% mitochondrial gene reads. 4553 HS1 cells were sequenced and 4355 were used in the analysis, 5674 PT1 cells were sequenced and 3620 were kept after filtering. The data was further log2-normalized and scaled to total expression in each cell, before further scaling cells on number of UMIs detected and percentage of mitochondrial gene count. PCA was run on variable genes defined using the FindVariableGenes function. tSNE was run using PCA dimensionality reduction.

### Proteomics

The cells were differentiated as previously described. Each line was divided into wells for differentiation in triplicates. At the day of harvest, the cells were washed twice with DPBS and dissociated using Accutase (25 μl/96 wells) for approximately 20 minutes until the cells had dissociated from the surface. The cells were then washed off with DPBS, centrifuged at 400 × g for 5 minutes and resuspended in 1 ml DPBS. The DPBS washings were repeated a total of three times. Finally, the supernatant was aspirated, and the pellets were snap-frozen on dry ice. The cells were later thawed and resuspended in 200 μl lysis buffer (50 mM DTT, 2w/v% SDS in 100 mM TRIS/HCl pH 8.6) and boiled at 95°C for 5 minutes, followed by vortexing and sonication (20 cycles of 15 seconds ON/OFF, 10 minutes in total; Bioruptor plus (model UCD-300, Diagenode)). Iodoacetamide was added to the final concentration of 100 mM in the samples and incubated at room temperature, in the dark for 20 minutes. The lysate was then clarified by centrifugation at 16000 × g for 5 minutes and the supernatant was collected. The proteins were precipitated using 9 volumes of cold EtOH and incubated overnight at −20°C. The samples were pelleted and washed once with 90% ethanol and dried with speed-vac. 100-120 μl 50 mM AmBic with 0.5% sodium deoxycholate was added to each sample and the samples were then sonicated for 15 minutes. 2 μg Sequencing Grade Modified Trypsin (Promega, Madison, WI) was added to each sample and digested overnight at 37°C. Sodium deoxycholate was removed from the samples by ethyl acetate extraction under acidic conditions. The peptide concentration was determined using the Pierce Quantitative colorimetric peptide assay (Thermo Fisher). The peptide concentration of the samples was adjusted to 140 μg/ml and analyzed using a Q Exactive HF-X mass spectrometer coupled to an Ultimate 3000 RSCLnano pump (Thermo Scientific). The peptides were loaded on an Acclaim PepMap100 C18 (5 μm, 100 Å, 75 μm i.d. × 2 cm, nanoViper) trap column and separated on an EASY-spray RSLC C18 (2 μm, 100 Å, 75 μm i.d. × 25 cm) analytical column. 0.1% formic acid (FA) was used for solvent A, and 80% acetonitrile (ACN) with 0.08% FA for solvent B. For peptide separation, a 120 min non-linear gradient was utilized with a flow-rate of 0.3 μl/min and the column temperature was set to 45°C. A top 20 data-dependent acquisition (DDA) method was applied, where MS1 scans were acquired with a resolution of 120,000 (@ 200 m/z) using a mass range of 375-1500 m/z, the target AGC value was set to 3e06, and the maximum injection time (IT) was 100 ms. The 20 most intense peaks were fragmented with a normalized collision energy (NCE) of 28. MS2 scans were acquired at a resolution of 15000, a target AGC value of 1e05, and a maximum IT of 50 ms. The ion selection threshold was set to 8.00e03 and the dynamic exclusion was 40 s, while single charged ions were excluded from the analysis. The precursor isolation window was set to 1.2 Th.

The raw files were analyzed with SEQUEST HT as a search engine using Proteome Discoverer (PD) v2.2 (Thermo Scientific). Human and chimpanzee samples were compared to the UniProt reference proteomes of human (UP000005640, downloaded on 2018-04-20, 71349 sequences), and chimpanzee (UP000002277, downloaded on 2018-04-20, 48770 sequences), respectively. The following search parameters were used: max. 2 missed cleavages, 10 ppm precursor, and 0.02 Da fragment mass tolerance, cysteine carbamidomethylation as a static modification, methionine oxidation, phosphorylation at serine, threonine, and tyrosine as dynamic modifications, acetylation as a dynamic N-terminal protein modification.

Each sample was run on two MS replicates, which were averaged for downstream analysis. Only proteins detected in at least one cell line with two valid values, and at least two biological replicates in both species were included in analysis. Additionally, 1% false discovery rate (FDR) at both peptide and protein levels was applied to include only high-confidence proteins. The resulting human and chimpanzee protein lists were then matched on gene names, resulting in 4971 proteins for analysis.

The protein intensity values were log2 transformed, scaled by subtracting the median of the sample, and missing data were imputed using Perseus v1.6.1.2. The R bioconductor package limma was used to fit a linear model and to compute moderated t-statistics [34].

### Statistical Analysis

To test if fold changes were significantly shifted from the main gene population, we performed permutation test (Fig 3f). For each group, 10000 random permutations were performed, selecting a random set of genes (n=#genes in original group). The difference in mean log2 fold change between the randomly selected gene set and all other genes in the data was calculated. We calculated the p-value as the frequency of randomly selected gene sets differing more from other genes than in the observed data.

For detection of differentially expressed genes at d14, we tested two cell lines/individuals per species. For RNA-seq data, each cell line was differentiated in two replicates, leaving 4 samples per species for differential expression analysis. DESeq2 [33] was used for normalization, dispersion estimates, model fitting and testing, and P < 0.001 (Benjamini-Hochberg adjusted), Wald’s test was used as significance cut-off. For proteomics, all four cell lines were differentiated in three replicates, and each differentiation was run on two separate mass-spectrometry runs. MS-replicates were pooled for final analysis, leaving 6 samples per species for differential expression analysis. lmFit and eBayes of limma [34] was used to fit a linear model and compute moderated t-statistics by empirical Bayes moderation of variance, with moderated p-value < 0.001 as a cut-off for significant differential expression.

## Acknowledgements

We would like to thank Stefan Thor and Volker Busskamp for comments on the manuscript and Fred H. Gage, Svante Pääbo, and Wieland Huttner for providing the chimpanzee iPSC lines. We are grateful to all members of the Jakobsson lab. We also thank J. Johansson, M. Persson Vejgården, U. Jarl, and A. Hammarberg for technical assistance. The work was supported by grants from the Swedish Research Council, the Swedish Foundation for Strategic Research, the Swedish Brain Foundation, the Swedish excellence project Basal Ganglia Disorders Linnaeus Consortium (Bagadilico), and the Swedish Government Initiative for Strategic Research Areas (MultiPark & StemTherapy). Support from the Swedish National Infrastructure for Biological Mass Spectrometry (BioMS) and ThermoFisher Scientific, San Jose USA is gratefully acknowledged.

## Author Contributions

DAG – investigation, supporting formal analysis, visualization, writing – original draft and review & editing. PLB – data curation, formal analysis, writing – original draft and review & editing. JGV – formal analysis, investigation, writing – review & editing. MR – resources, writing – review & editing. MEJ – visualization, writing – review & editing. SN– methodology, writing – review & editing. MP – methodology, funding acquisition, writing – review & editing. GMV – resources, writing – review & editing. JJ – conceptualization, funding acquisition, project administration, writing – original draft and review & editing.

## Author Information

The authors declare no competing interests.

## Code availability

The code is available at https://github.com/perllb/HsPt_Protein-RNA

---

**Supplementary Fig. 1.**
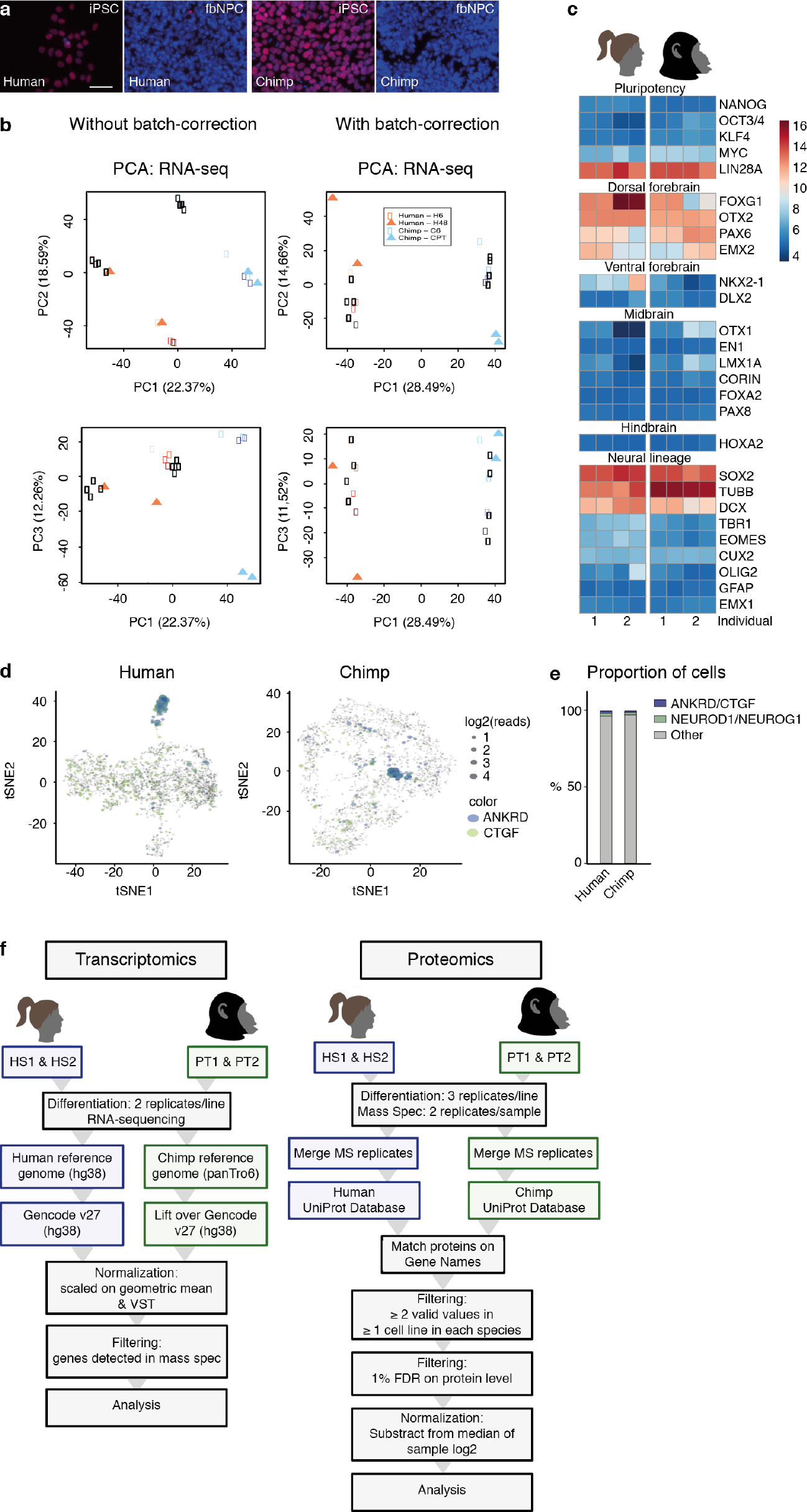
(A) Immunocytochemical staining of NANOG (red) in human and chimpanzee fbNPC and iPSC, DAPI (blue) is used as counterstain. The scale bar represents 50 μm. (B) Principal component analysis plots of RNA-seq data from human and chimpanzee, before and after batch correction. (C) Heatmap of neural marker transcript expression at day 14 of differentiation. (D) tSNE of human and chimpanzee single-cell RNA-seq data showing the expression of endothelial markers ANKRD1 and CTGF. (E) Bar charts showing the percentage of ANKRD1/CTGF+ as well as NEUROD1/NEUROG1+ cells in single-cell RNA-seq. (F) Schematic figure illustrating an overview of samples used in the RNA-seq and mass spectrometry experiments, and the analysis strategy for both data sets.

**Supplementary Fig. 2.**
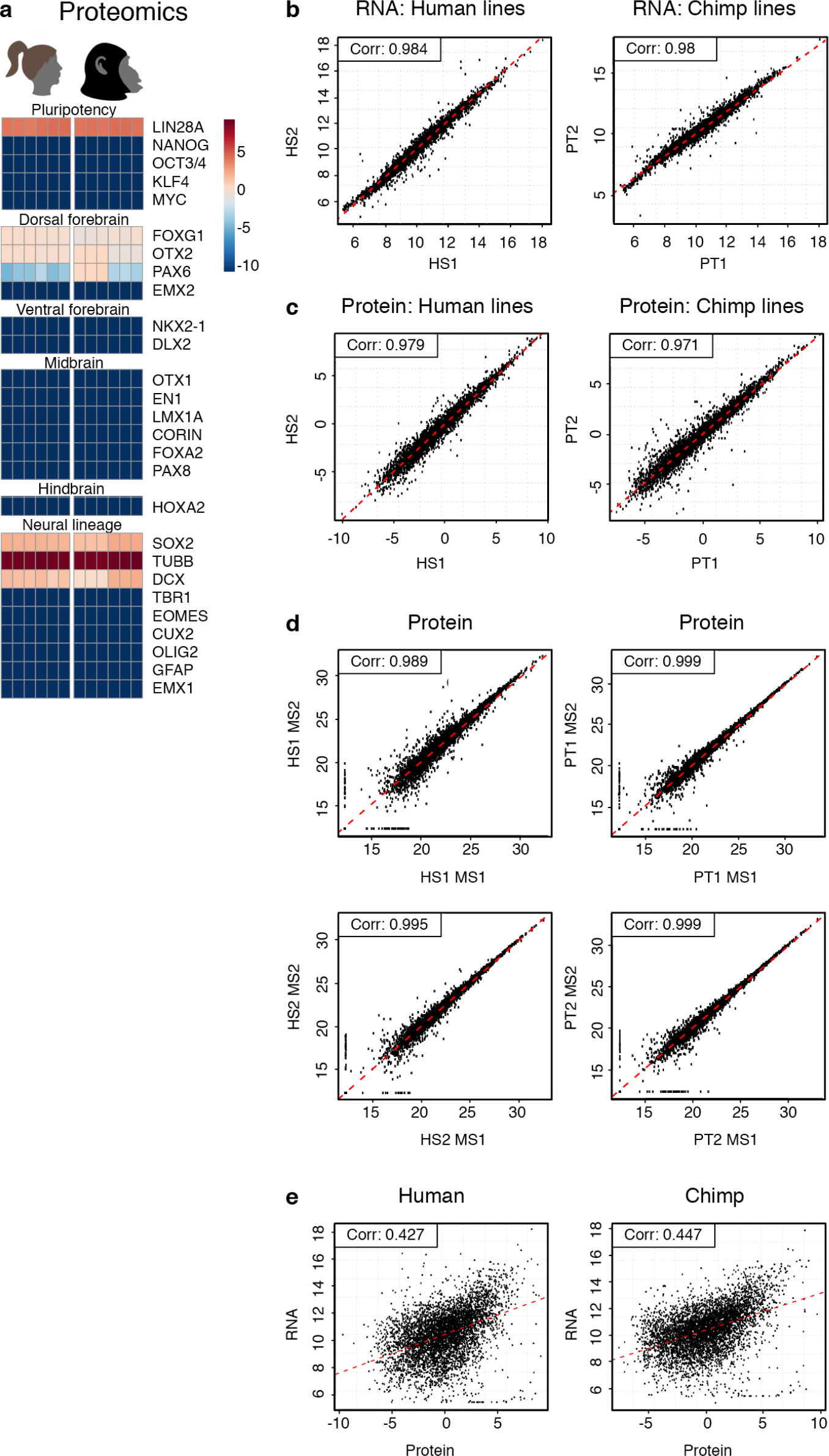
(A) Protein marker characterization of fbNPCs. (B) Abundance correlation comparing RNA levels between individuals of the same species: Pearson corr. human = 0.984, chimpanzee = 0.98. (C) Abundance correlations of protein levels between individuals of the same species: Pearson corr. human = 0.979; chimpanzee = 0.971. (D) Protein abundance correlation between the two MS replicates for each line. (E) Correlation between RNA and protein levels for human (Pearson corr. = 0.427) and chimpanzee (Pearson corr. = 0.447).

**Supplementary Fig. 3.**
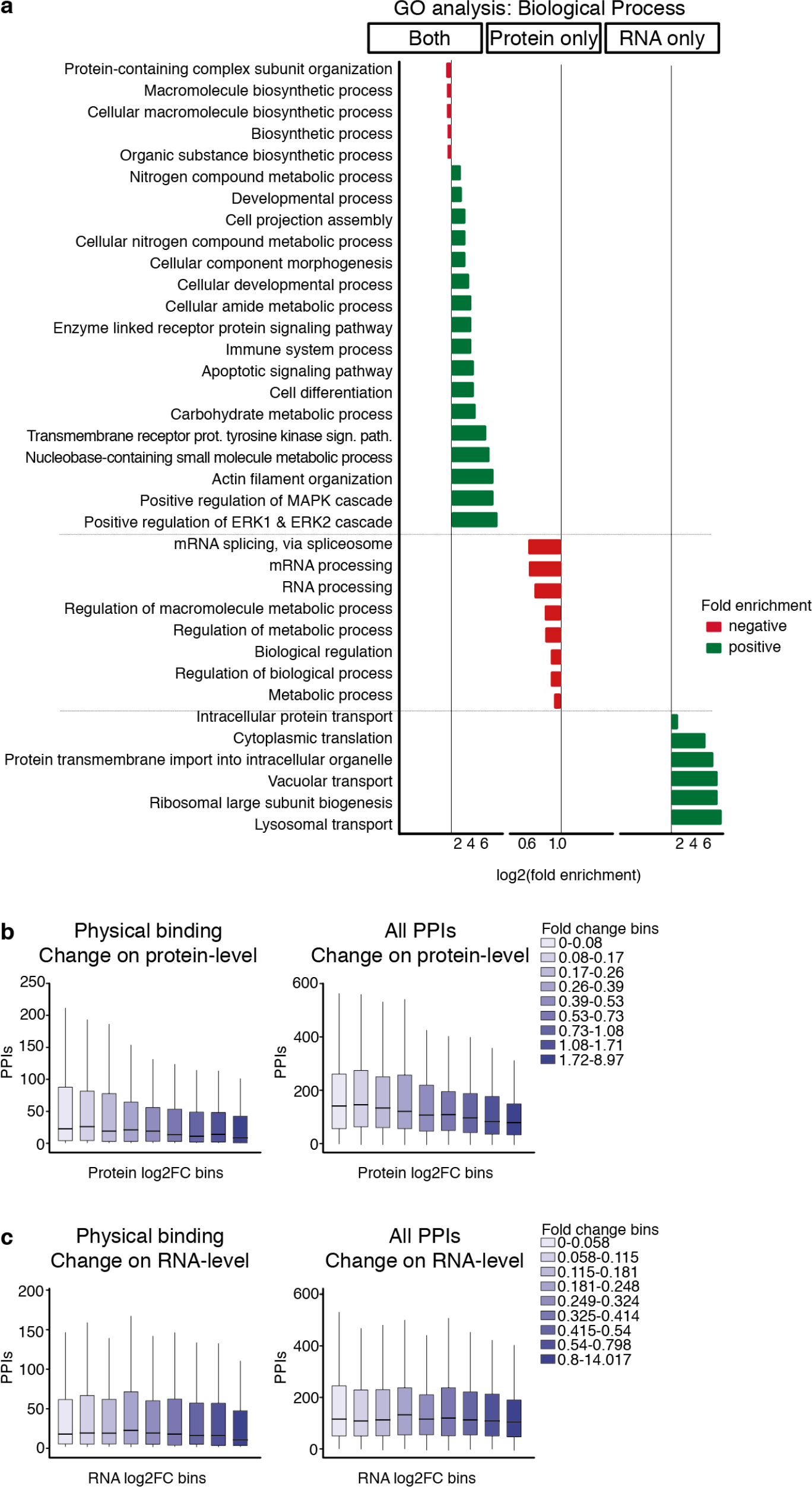
(A) Gene ontology (GO) analysis of biological process for differentially expressed genes. (B) Boxplots showing the number of PPIs: physical binding to other proteins only, and all PPIs. The proteins were divided into 9 subgroups with equal numbers of proteins per group, based on the fold change between human and chimp. The lower and upper hinges correspond to the first and third quartiles. The upper whisker extends from the hinge to the highest value, no further than 1.5 × IQR from the hinge (where IQR is the inter-quartile range, or distance between the first and third quartiles). The lower whisker extends from the hinge to the lowest value, at most 1.5 × IQR of the hinge. (C) The same analysis as in B, based on human-chimpanzee RNA divergence.

